# Roles Played by Enhancer of Split Transcription Factors in *Drosophila* R7 Photoreceptor Specification

**DOI:** 10.1101/2025.01.30.635716

**Authors:** A. Arias Ronald, E. Mavromatakis Yannis, Andrew Tomlinson

## Abstract

When a cell receives multiple developmental signals simultaneously, the intracellular transduction pathways triggered by those signals are coincidentally active. How then, do the cells decode the information contained within those multiple active pathways to derive a precise developmental directive? The specification of the Drosophila R7 photoreceptor is classic model system for investigating such questions. The R7 fate is specified by the combined actions of the of the Notch (N) and receptor tyrosine kinase (RTK) signaling pathways. The two pathways cross-communicate in an integrative mechanism but also supply information independently of each other. Collectively, this information is summed to provide an unambiguous directive for the R7 fate. Our goal is to understand these mechanisms. Here, we examine how N activity represses transcription of the *phyllopod* gene in the process of information integration with the RTK pathway, and how it represses expression of the *seven-up* gene in an independent mechanism needed for R7 fate. We describe how N activity achieves these transcriptional repressions, and identify Enhancer of Split transcription factors as the mediators of those actions.

**Summary Statement:** This work examines the role played by Enhancer of Split transcription factors in mediating N signals in Drosophila R7 photoreceptor specification.

## Introduction

The last decades have seen great success in our understanding of the many signaling pathways that direct cell fates. The receptors and ligands have been identified as have the intra cellular molecular transduction pathways that they activate. What remains unclear, is what happens in cells that receive multiple developmental signals simultaneously. How is the information encoded in those multiple active pathways decoded to derive a precise fate directive? The transduction pathways typically regulate gene expressions, but when different signalings occur simultaneously, the resulting fate is rarely specified by the simple summation of the outputs of the individual transduction pathways. Rather, molecular integration of the two pathways results in a novel transcriptional profile which specifies the fate. The *Drosophila* R7 photoreceptor is specified by the combined actions of the receptor tyrosine kinase (RTK) and Notch (N) signaling pathways^1,2^, and we use this as a model system to understand how the information present in the two pathways is decoded to provide the R7 fate directive. In this process, the R7 precursor derives two sequential pieces of information. First, in an integrative interaction between the two pathways it is specified as a photoreceptor precursor (rather than a non-photoreceptor cone cell). Second, in a process that appears to be directed only by N signaling, it is directed to adopt the R7 rather than the R1/6 photoreceptor fate^3^. Here, we investigate the molecular mechanisms that underlie these two developmental steps.

The R7 precursor is derived from a pool of cells that also gives rise to the R1/6 photoreceptors and the lens-accessory cone cells. The cells of this pool express the Drosophila EGF Receptor (DER – the general RTK employed in *Drosophila* eye development) and N on their plasma membranes, and the repressive transcription factor Tramtrack (Ttk) in their nuclei (Fig.1E)^4^. Ttk blocks the specification of the photoreceptor fate, and the combined actions of the RTK and N signaling pathways direct its removal in photoreceptor precursors and its persistence in presumptive cone cells. Ttk removal is controlled by the transcription of the *phyllopod* (*phyl*) gene, which encodes an adaptor protein that organizes the assembly of a Ttk degradation complex^5–9^. *phyl* transcription is regulated by two ETS transcription factors both regulated by the RTK pathway. One is Yan, a transcriptional repressor of *phyl* transcription and the other is PointedP2 (PntP2) which actively promotes the gene’s expression^10–12^. Upon DER activation, MAPK translocates to the nucleus where it phosphorylates both Yan and PntP2; inactivating the former and potentiating the latter (Fig.1F). *phyl* transcription is active in all cells of the developing eye^3^, and in photoreceptor precursors, the modulations of Yan and Pntp2 activities increase its expression levels which promotes the removal of Ttk. This increase in *phyl* transcription thus acts as a switch; when it occurs, Ttk is removed and the photoreceptor fate is specified, when not, Ttk persists and the non-photoreceptor fate is imposed.

**Figure 1.**
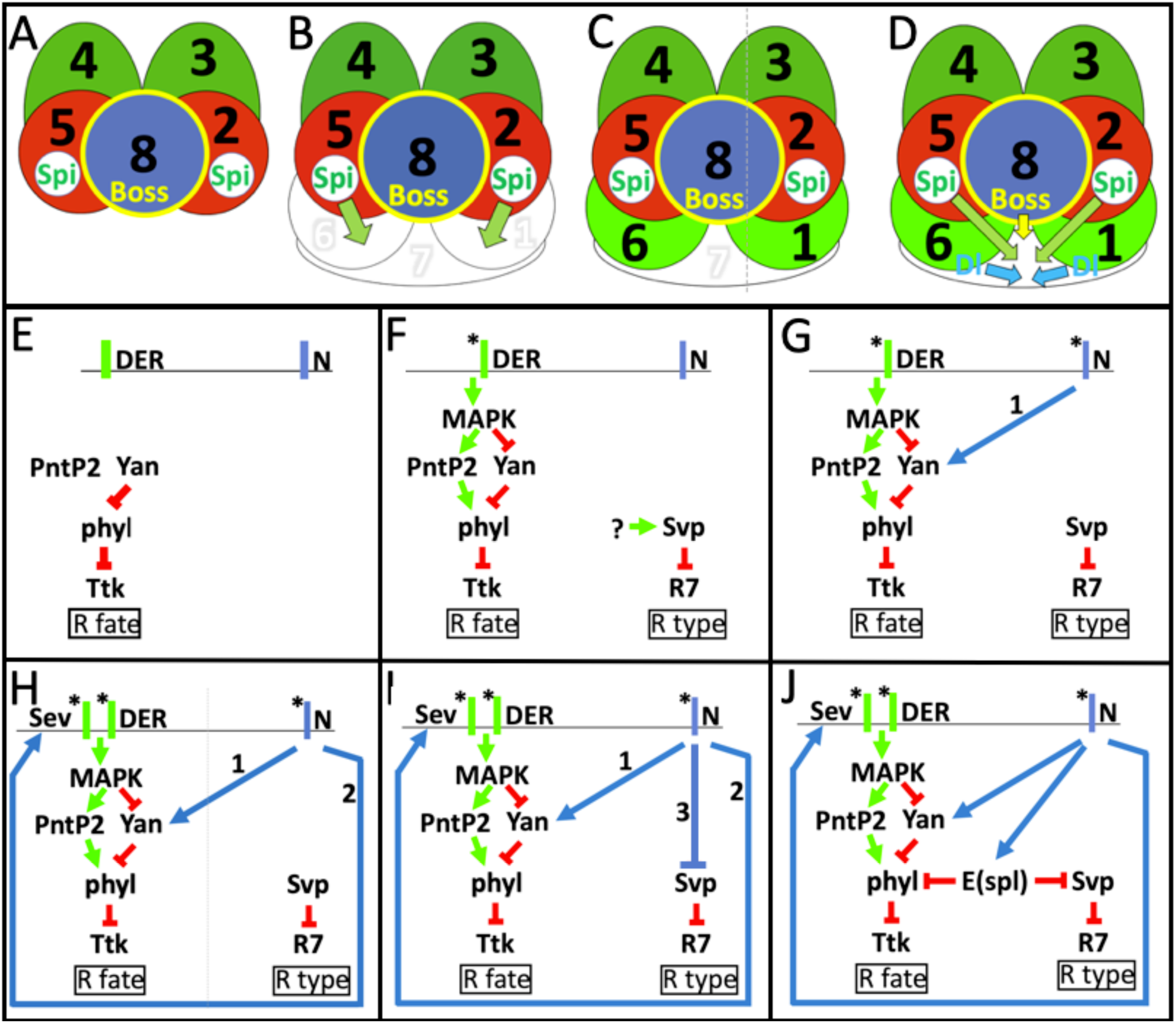
Schematic descriptions of the specifications of the R7 and R1/6 photoreceptors. (A-D) The recruitment of the R1/6/7 precursors and signals the signals that specify them. (A) The precluster is made from the presumptive R2,3,4,5,8 photoreceptors. R2/5 release Spitz, the DER ligand, and R8 expresses the Sev ligand Boss on its membrane. (B) Three naïve cells (white) are recruited to the R2/8/5 face of the precluster. R1/6 contact the Spitz-expressing R2/5 which activates their DER. The middle cell (R7) does not immediately receive Spitz. (C) The activation of DER specifies R1/6 types (green) while the R7 precursor waits. (D) R1/6 express Dl which activates N in R7 and provides Sev which is activated by Boss on R8. When Spitz reaches it, N, DER and Sev are active in the R7 precursor and collectively specify R7’s fate. (E) The naïve cells express DER and N on their membranes, and Yan inhibits *phyl* transcription, resulting in TTK persistence, and the non-photoreceptor fate. (F) In R1/6 precursors, the DER pathway inactivates Yan and promotes Pntp2, resulting in *phyl,* transcription and Ttk removal. The R1/6 precursors also express Svp which represses the R7 fate. It is unclear what promotes Svp expression. (G) N activity promotes (1) *yan* transcription increasing Yan protein levels that titrate MAPK’s efforts to inactivate it; *phyl* levels are not raised, Ttk persists and opposes photoreceptor specification. (H) N activity promotes (2) *sev* transcription which supplies the Sev RTK to the cell. Sev hyperactivates MAPK, overcoming the Yan opposition; Ttk is removed and photoreceptor specification occurs. (I) N activity prevents (3) Svp expression and the R7 fate-block. (J) N activates *E(spl)* transcriptions which repress the expression of the *phyl* and *svp* genes.

In R1/6 precursors, in which N is inactive, the switch occurs in a relatively simple manner (Fig.1F), but in presumptive R7s, N activity both opposes and promotes *phyl* transcription and the switch occurs in a more complicated way. To oppose *phyl* transcription, N activity hyperactivates *yan* transcription and the resulting increase in Yan protein levels is thought to counter the ability of MAPK to inactivate it (Fig.1G)^3^. To promote *phyl* transcription, N activity drives *sev* transcription that supplies high levels of the Sevenless (Sev) RTK to the cell (Fig.1H)^13^. The activation of Sev hyperactivates the RTK transduction pathway and overcomes the increased amounts of Yan, and the increase in *phyl* transcription ensues. Thus, although N activity in the R7 precursor triggers a molecular “dance” within the cell, the output is the same as in R1/6 precursors; the increased transcription of *phyl*, the resulting degradation of Ttk and the specification of the photoreceptor fate.

Once the R7 precursor is defined as prospective photoreceptor (the removal of Ttk), N activity then directs the specification of the R7 rather than R1/6 class (Fig.1I). This is achieved by preventing expression of the *seven-up (svp)* gene in the R7 precursor. Svp domineeringly imposes the R1/6 fate and its absence from the R7 precursor is necessary for its appropriate specification^3,14,15^.

The *Enhancer of split gene complex (E(spl))* includes seven classic N response genes that encode helix-loop-helix repressive transcription factors that are used repeatedly throughout development to suppress the transcription of their target genes^16,17^. Earlier work suggested that a member of this family (E(spl)mδ)^1^ was active in R7 specification as evidenced by its potency to redirect R1/6 precursors to the R7 fates. Extending this observation, we used ectopic/overexpression studies to evaluate the ability of each of the seven *E(spl)* genes to influence R7 specification and found that three (*E(spl)mδ/β/8*), when expressed at moderate levels, re-specified R1/6 precursors as R7s, and at higher levels directed the differentiation of non-photoreceptor cone cells. This indicated that E(spl) proteins could disturb both steps in R7 specification, both of which are controlled by N activity. We therefore generated transcriptional reporters for the three and found that *E(spl)*mδ and *E(spl)*m8 were transcribed in the R7 precursor, but to our surprise, *E(spl)mβ* was not. We infer that the potency *of E(spl)mβ* expression to change the fate of R1/6 precursors results from it mimicking the actions of other *E(spl)* genes. This invalidated ectopic/overexpression experiments as a method to identify the *E(spl)* genes active in R7 specification. We therefore switched to loss of function analyses and generated a deletion of the *E(spl)-complex*. Clones of this deletion revealed that *E(spl)* gene functions are required for the specification of the R7 fate, and are specifically required for the repression of *phyl* and *svp* transcriptions (Fig.1J).

## Results

### Expression of *E(spl)mδ, E(spl)m8* and *E(spl)mβ* in R1/6 precursors mimics N activation

Since E(spl) proteins typically mediate N signals and since *E(spl)mδ* had been implicated in the R7-v-R1/6 choice we investigated the roles of E(spl) transcription factors in R7 specification. We expressed *UAS* versions of the seven *E(spl)* genes using the *sev.Gal4* driver line. *sev* transcription is absent from R1/6 precursors, but fortuitously, *sev.Gal4* is active in R1/6 along with R7 and cone cell precursors. Of the seven, only *E(spl)mδ, E(spl)m8* and *E(spl)mβ* promoted fate changes in R1/6 precursors. We had no independent method of validating the *UAS* lines, and for those that did not disturb R1/6 fates, we do not conclude that these play no roles in R7 specification. Rather, we report that we did not detect phenotypes for whatever reason.

Ectopic N activation in R1/6 precursors (*sev.N\**) induces two phenotypes; the cells transform into R7 types (Fig. 2B). but in a *sev°* background, they become cone cells and usually four large-rhabdomere cells (R2,3,4,5) are evident in adult eyes (Fig. 2C)^2^. Both phenomena were phenocopied by the expressions of the three *E(spl)* genes. Adult eye sections showed the multiple R7 phenotype with the corresponding loss of large-rhabdomere R1/6 photoreceptors (Fig, 2D,E), and at higher transgene expressions, we observed ommatidia lacking R7s with a reduced complement of large-rhabdomere photoreceptors (Fig.2F), which occurs when R1/6/7 precursors become cone cells (Fig.2C). For example, when single copy UAS *E(spl)mδ* was present; of N = 227 ommatidia: 190 were wild type and 37 showed supernumerary R7s. But when two copies of *E(spl)mδ* were present, of N = 189 ommatidia: 4 were wild type and 185 showed ectopic R7s and/or the loss of photoreceptors. These stronger phenotypes also occurred when any of the three transgenes were jointly expressed with any other. For example, when *E(spl)mδ* was coexpressed with *E(spl)mβ*, of N =161 ommatidia:16 were wild type and 145 showed ectopic R7s and or the loss of photoreceptors. These results indicate that ectopic/over-expression of *E(spl)* genes mimics N activity in specifying presumptive R1/6 cells as R7 types and in directing R1/6/7 precursors to become cone cells.

**Figure 2.**
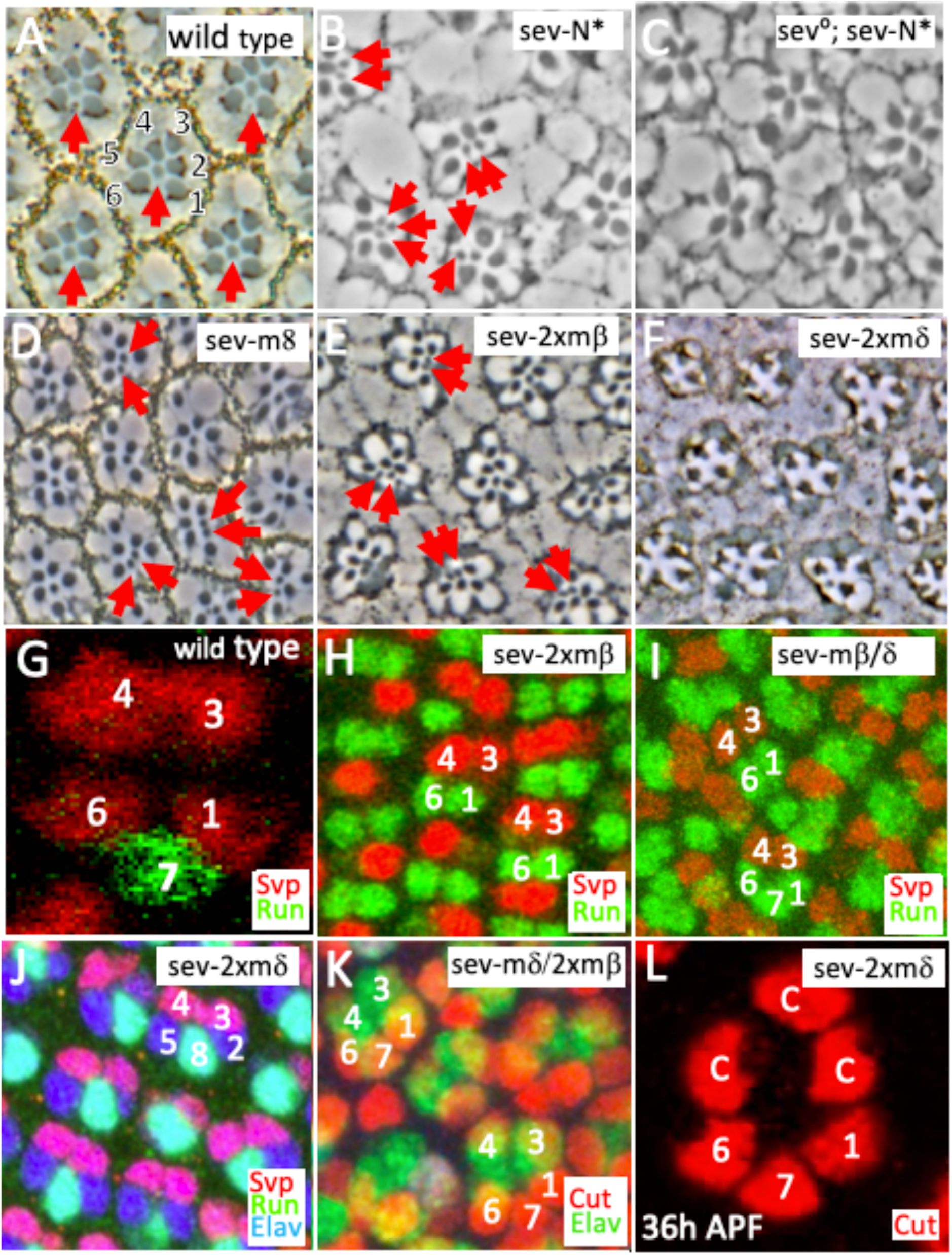
Expression of *E(spl)* genes in R1/6 precursors re-specifies them as R7s or cone cells. (A-F) Adult eye sections. (G-L) Larval/pupal eye tissues (A) In *wild type* ommatidium, a single small-rhabdomere R7 (red arrow) is surrounded by six large-rhabdomere photoreceptors. (B) In *sev.N\** eyes, N activity in R1/6 precursors can re-specify them as R7s (red arrows) with a corresponding reduction in the number of large-rhabdomere photoreceptors. (C) In *sev°*; *sev.N*,* the specification of R1/6/7 as cone cells leaves only four large-rhabdomere photoreceptors. (D,E) When *UAS.E(spl)m8* or two copies of *UAS.E(spl)mβ* transgenes are expressed by *sev.Gal4*, extra R7s (red arrows) are evident at the expense of large-rhabdomere types. (F) When two *UAS.E(spl)mδ* transgenes are expressed by *sev.Gal4*, ommatidia lacking R7s with reduced numbers of large rhabdomere cells are evident mimicking the effects of transformation of R1/6/7 precursors into cone cells. (G) In *wild type* clusters, R1/6/3/4 precursors express Svp (red) and the presumptive R7 expresses Runt (green). (H,I) When two *E(spl)mβ* transgenes, or when one copy of *E(spl)mβ* is jointly expressed with *UAS.E(spl)mδ,* cells in the R1/6 positions frequently appear as R7s evidenced by their Runt label (green) and not Svp (red). (J) When two *UAS.E(spl)mδ* transgenes are expressed by *sev.Gal*, clusters containing only precluster photoreceptors showing varied Elav (blue), Svp (red) and Runt (green). (K) When a *UAS.E(spl)mδ* and two *UAS.E(spl)mβ* transgenes are expressed by *sev.Gal4,* R1/6 precursors transform into cone cells evidenced by their expression of Cut (red) and absent Elav staining (blue). (L) In a 36h pupa retina in which *two UAS.E(spl)mδ* transgenes are expressed by *sev.Gal4,* five cone cells are frequently observed as indicated by their Cut expression (red). From their positions, three appear to be derived from R1/6/7 precursors.

To assess the validity of these fate transformations, we examine eye discs in which the *E(spl)* transgenes were expressed as in the manner above. The transformation of R1/6 precursors into either R7 or cone cell types was evident. Fig. 2H,I show the effects of expression of two copies of *E(spl)mδ,* or a single copy jointly expressed with *E(spl)mβ.* In each, R1/6 precursors transform into R7s as evidenced by their expression of the R7 marker Runt at the expense of the R1/6 label Svp. Transformation of R1/6/7 precursors into cone cells was evidenced by their loss of photoreceptor markers (Fig. 2J) and by the gain of the cone cell label Cut (Fig. 2K). To corroborate the transformation of R1/6/7 into cone cells, we examined 36h pupal eyes when two copies of *E(spl)mδ* were expressed and observed cone cell groupings made frequently from 5 or 6 cells (rather than the normal 4), with three arranged in the R1/6/7 characteristic arrangement (Fig.2L). This suggests that following the reassignment of R1/6/7 precursors as cone cells, the ommatidia recruit two or three more cells to cone cell positions.

These results suggest that higher levels of *E(spl)* gene expressions mimic N functions that opposes the actions of the RTK pathway to specify the photoreceptor fate, and that at lower levels, it phenocopies N roles that direct the R7 rather than R1/6 fate. Since the three transgenes occupy different genomic locations and we had no method for assessing their expression levels, we do not suggest that any is natively more potent than the others in its abilities to change cell fates.

### Generation of *E(spl)m*δ, *E(spl)m8 and E(spl)mβ* transcriptional reporters

If the three *E(spl)* genes relay N information in the R7 precursor, then they should be expressed therein. To test this, we generated transcriptional reporters for the three using Crispr-induced homologous repair in which the coding sequence of a fluorescent protein with a nuclear localization signal was inserted into the first coding exon of the genes using the native ATG as the initiation codon. These reporters were also null mutations of the genes into which they were inserted

### The expression pattern of *E(spl)mδ.GFP*

Figure 3 shows the expression pattern of *E(spl)mδ.GFP^NLS^* in the eye-antennal, wing and leg discs (A,B,C, respectively). In eye discs, GFP expression was detected at low levels in the majority of cells posterior to the furrow (Fig.3D), and was upregulated in R3/4, R7 and the cone cell precursors (Fig.3E-H) before fading to moderate levels. R3/4, R7 and cone cell precursors are all N responsive, suggesting that *E(spl)mδ* transcription is likely controlled by N activity. This expression pattern differs substantially from that of the *mδ.0.5-lacZ* reporter previously thought to recapitulate mδ transcription^1^. *E(spl)mδ.GFP^NLS^* null homozygous animals were fertile and healthy. In sections through adult eyes (not shown), of N = 313 ommatidia, 311 were wild type and two showed 7 large rhabdomere photoreceptors. Thus, the loss of *E(spl)mδ* function produces almost completely wild type eyes with no disruption of R7 specification.

**Figure 3.**
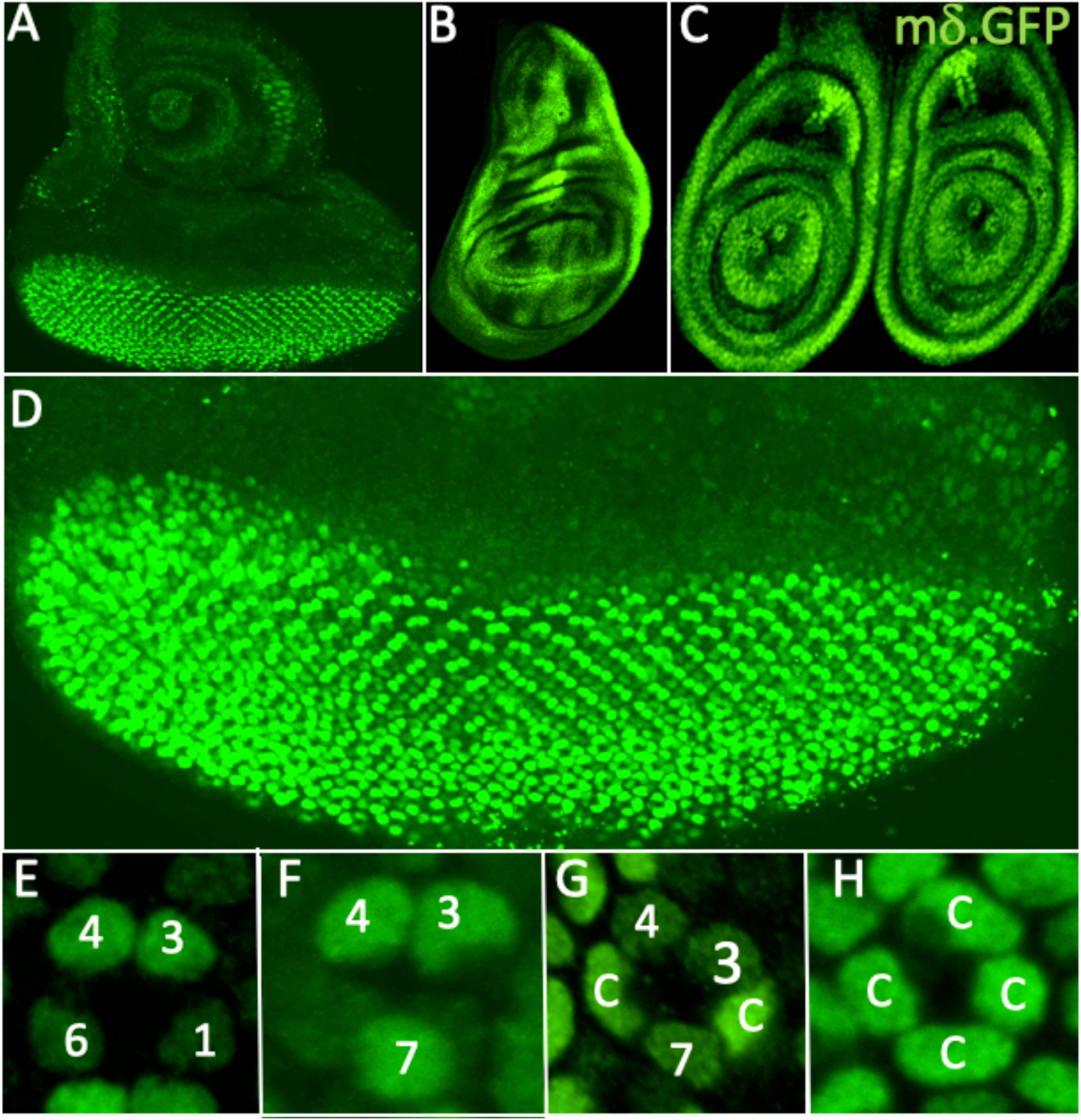
Expression patterns of *E(spl)mδ.GFP^NLS^* in imaginal discs. (A-C) expression in (A) eye-antennal (B) wing and (C) leg discs. (D) In eye discs, staining is evident in cells posterior to the furrow. (E) Staining is initially strong in R3/4 cells. (F) R7 stains strongly. (G) R3/4 and R7 staining reduces as strong expression occurs in two cone cell precursors (c). (H) The group of four cone cells all stain strongly.

### The expression pattern of *E(spl)m8.GFP*

Figure 4 shows the expression pattern of *E(spl)m8.GFP^NLS^* in the eye-antennal, wing and leg discs (A,B,C, respectively). In eye discs, GFP expression was detected in the majority of the cells posterior to the furrow (Fig.4D), and was upregulated in R3/4 precursors (Fig.4E) followed by a strengthening in R4 as that of R3 declined (Fig.4F). Moderate staining was then observed in R1/6 precursors and more strongly in the presumptive R7 (Fig.4F) and cone cell precursors (Fig4.G,H). Strong N activation occurs in R4 precursors^18–20^ which comports with *E(spl)m8* mediating its actions here. Furthermore, the expression in R7 and cone cell precursors corresponds with N activation in these cells. *E(spl)m8.GFP^NLS^* null homozygotes were fertile and healthy, and sections through adult eyes (not shown) showed almost completely wild type ommatidia; of N = 214 ommatidia scored: 204 were normal, 8 showed failures to resolve R3/4 fates, and 2 were disrupted in a manner that cannot be easily classified. Thus, loss of *E(spl)m8* gene function produced almost completely wild type eyes with no evident effects on R7 specification.

**Figure 4.**
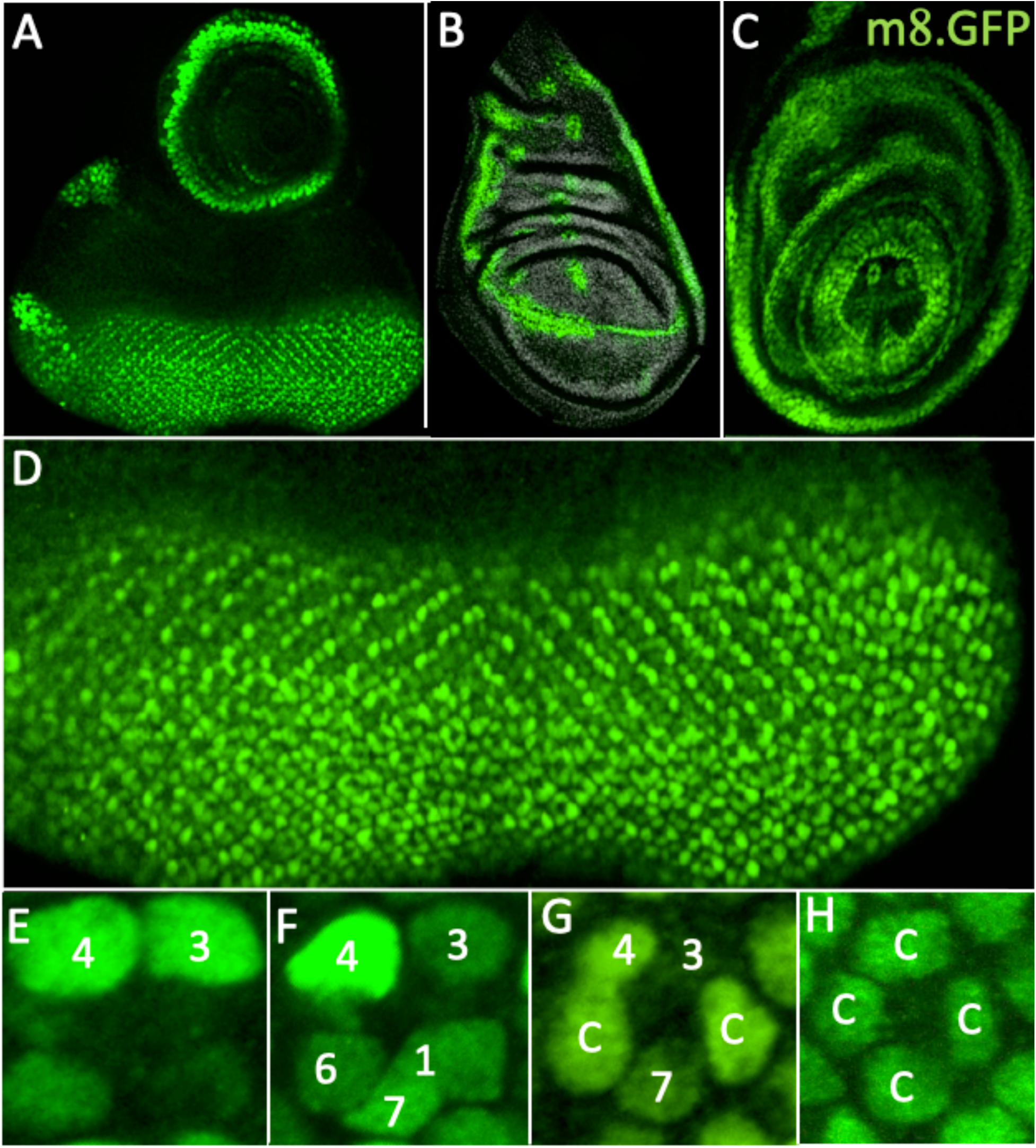
Expression patterns of *E(spl)m8.GFP ^NLS^* in imaginal discs. (A-C) expression in (A) eye-antennal (B) wing and (C) leg discs. (D) In the eye disc, staining is evident in cells posterior to the furrow. (E) Staining is initially strong in R3/4 cells. (F) R4 staining strengthens as R3 weakens. (F) R7 Staining is higher than R1/6. (G) As the staining in R7 weakens, two cone cell precursors (c) stain more strongly. (H) The four cone cell precursors stain at moderate levels.

### The expression pattern of *E(spl)mβ.BFP*

Figure 5 shows the expression pattern of *E(spl)mβ.GFP^NLS^* in the eye-antennal, wing and leg discs (A,B,C, respectively). We changed the reporter from GFP to BFP here to increase our repertoire of fluorophores. In eye discs, expression began weakly in R3/4 precursors followed by a strong upregulation in R4 (Fig.5E) as defined by its position in the quartet of R1/6/3/4 precursors highlighted by Svp expression (red, Fig.5E’). There is no staining of R7 precursors indicating that *E(spl)mβ* is not transcribed therein and any effects the gene’s expression engenders do relate to a native role in R7 specification but likely result from it mimicking the functions of other E(spl) family members. This result invalidated the use of expression studies to identify the *E(spl)* genes active in R7 specification. We therefore turned to loss of function experiments.

**Figure 5.**
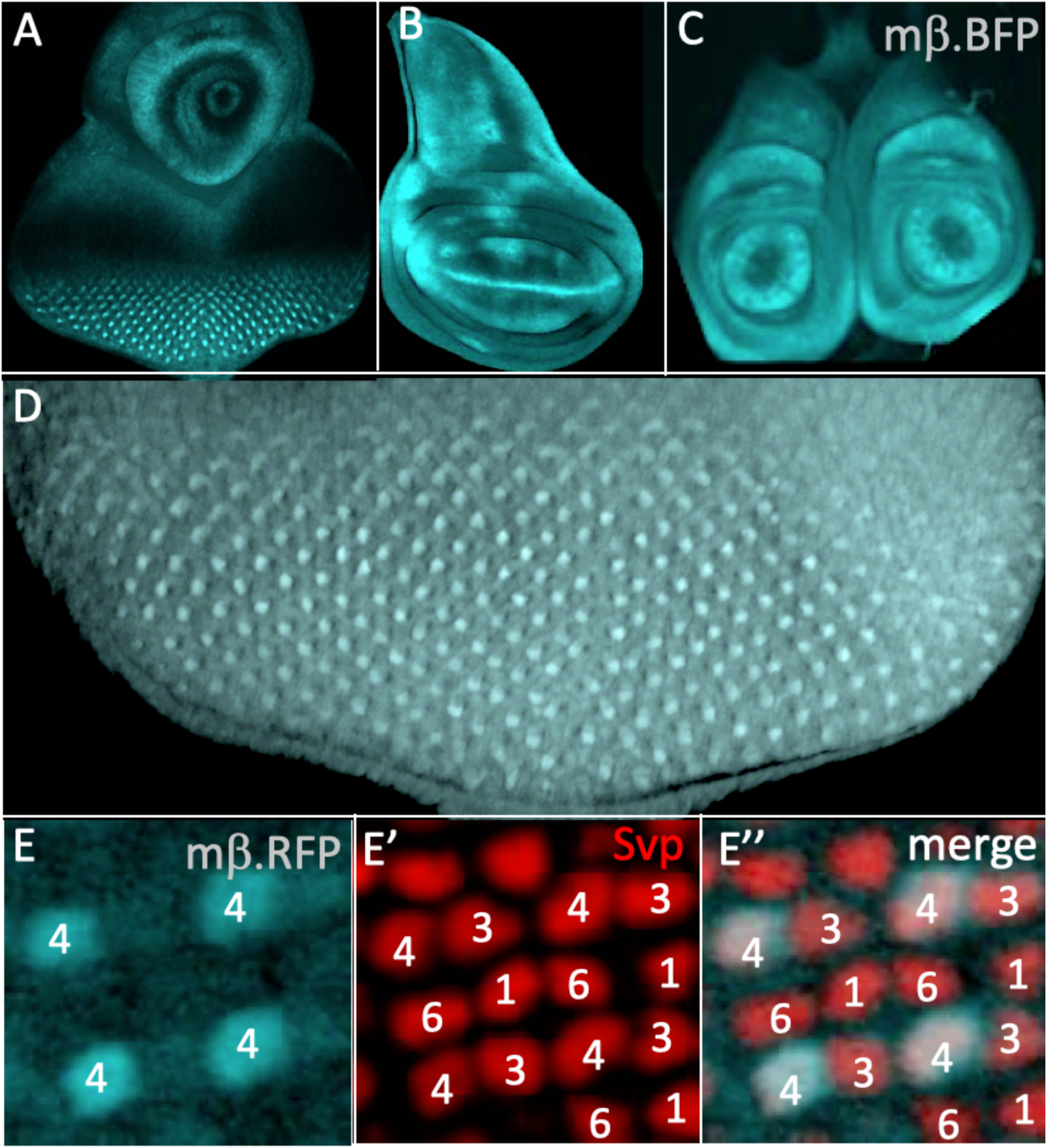
Expression patterns of *E(spl)mβ.BFP ^NLS^* in imaginal discs. (A-C) expression in (A) eye-antennal (B) wing and (C) leg discs. (D) In the eye disc, strong staining is evident in cells posterior to the furrow. (E) Staining is seen in R4 precursors as evidenced by R1/6/3/4 highlighted by Svp (red).

### The *E(spl)mδ* and *E(spl)m8* double mutant

Since *E(spl)mδ* and *E(spl)m8* both influenced cell fate specifications and were expressed in the R7 precursor, we generated a double-mutant chromosome to evaluate their joint loss of function phenotypes. Onto a previously generated an *E(spl)m8.mCherry* chromosome, we inserted *E(spl)mδ.GFP^NLS^* generating the double mutant (Fig.6A). Homozygous animals were healthy and fertile, and sections through adult eyes showed that the vast majority of ommatidia appeared normal; of N = 255: 240 were wild type, 7 showed defects in R3/4 position two had 7 large-rhabdomere photoreceptors, and one lacked an R7. The yellow circle in Fig.6B indicates an ommatidium with seven large-rhabdomere photoreceptors, and the red circle one in which R3/4 do not adopt their correct positions. Since both *E(spl)m8* and *E(spl)mδ* are expressed in the R7 precursor and their combined mutant phenotypes showed largely wild type ommatidia, this suggested that redundant functions were encoded in other genes. To determine whether these were also *E(spl)*genes, we generated a deletion of the complex and examined its phenotypes

**Figure 6.**
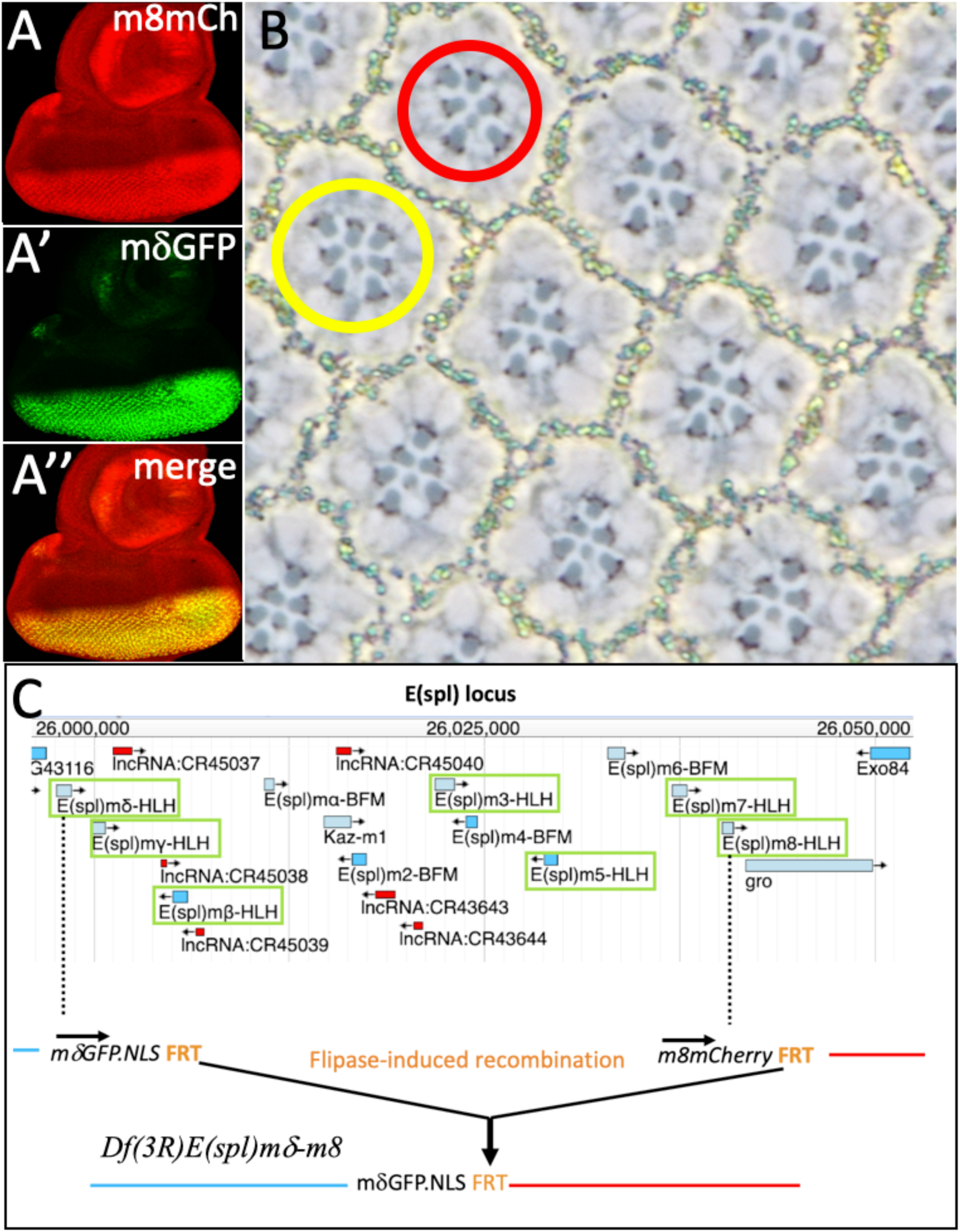
*E(spl)* mutant phenotypes and the generation of an *E(spl)-complex* deficiency. Onto an *E(spl)m8.mCherry* chromosome (A), *E(spl)mδ.GFP^NLS^* was inserted (A’), generating a chromosome bearing both reporters (A’’). (B) Section of an adult retina of a homozygous *E(spl)mδ.GFP^NLS^ E(spl)m8.mCherry* double mutant showing mostly wild type ommatidia. The red circle indicates an ommatidium with R3/4 mispositioned, and the yellow circle highlights an ommatidium with 7 large-rhabdomere photoreceptors. (C) Schematic details of the generation of the *Df(3R)E(spl)mδ-m8* chromosome, adapted from a FlyBase JBrowse image of the *E(spl)* locus. The complex contains seven genes encoding transcriptional repressors (green boxes). The arrows indicate directions of transcription. *E(spl)mδ* and *E(spl)m8* lie at the proximal and distal extremes of the complex. They are transcribed in the same direction. Each has a 3’FRT. Heat-shock induced Flipase expression the sequences between the FRTs to generate the deficiency.

### A deletion of *the E(spl)* complex

Fortuitously, the *E(spl)*m8 and *E(spl)*mδ genes lie at the proximal and distal ends of the complex and their directions of transcriptions are the same (Fig.6C). Both *E(spl)mδ.GFP^NLS^* and *E(spl)m8.mCherry* transgenes carry 3’ FRTs, and Flipase expression was used to induce recombination between them to generate a functional deletion of the complex (Fig.6C). The resulting *Df(3R)E(spl)mδ−m8* chromosome retained the *E(spl)*mδ.GFP^NLS^ reporter (an *E(spl)mδ* null allele) followed by the FRT and the sequences of the mutated *E(spl)m8* gene. The expression pattern of *mδ.GFP^NLS^* on chromosome was modified from the original, likely resulting from the loss of *E(spl)*mδ enhancers or the de novo proximity of those of local to *E(spl)m8*.

### Clonal phenotype of the loss of the *E(spl)-complex*

We recombined *FRT82* onto the *Df(3R)E(spl)mδ-m8* chromosome and generated clones in adult eyes marked by *w°* (the absence of pigment) by providing heat-shock-induced Flipase at different times of development to produce mosaic patches of varying sizes. With the exception of small late clones, the majority produced scars in the eyes that prevented any analysis of the requirement for the *E(spl)* genes in R7 specification. We therefore used *GMR.flip* (which provides Flipase expression after the preclusters have formed) to generate clones in the R1/6/7 and cone cell precursor populations. Of 46 normally constructed ommatidia that showed mosaicism within the R1/6/7 triumvirate, all 46 had a pigmented (wild type) R7s while 30 R6s and 18 R1s were w^-^ (mutant). Thus, although the *E(spl)-complex* could be removed form R1/6s without affecting their specification, its presence was critically required for R7 assignment. Furthermore, w^-^ (mutant) cells in the R7 positions appeared as large-rhabdomere R1/6 types (black arrows, Fig.7A), suggesting that in the absence of the *E(spl)-complex*, R7 precursors are specified as R1/6 types. Additionally, supernumerary large-rhabdomere photoreceptors were observed in mosaic ommatidia (red arrow, Fig.7A inset) likely resulting from mutant cone cell precursors transforming into R1/6 types in the manner that occurs when these cells lack N function^13^. This suggests that loss of the *E(spl)-complex* results in the loss of the N-induced opposition to photoreceptor specification (Fig.1G) and (along with the loss of the R7-specifying information) results in mutant cone cell precursors being specified as R1/6 types.

**Figure 7.**
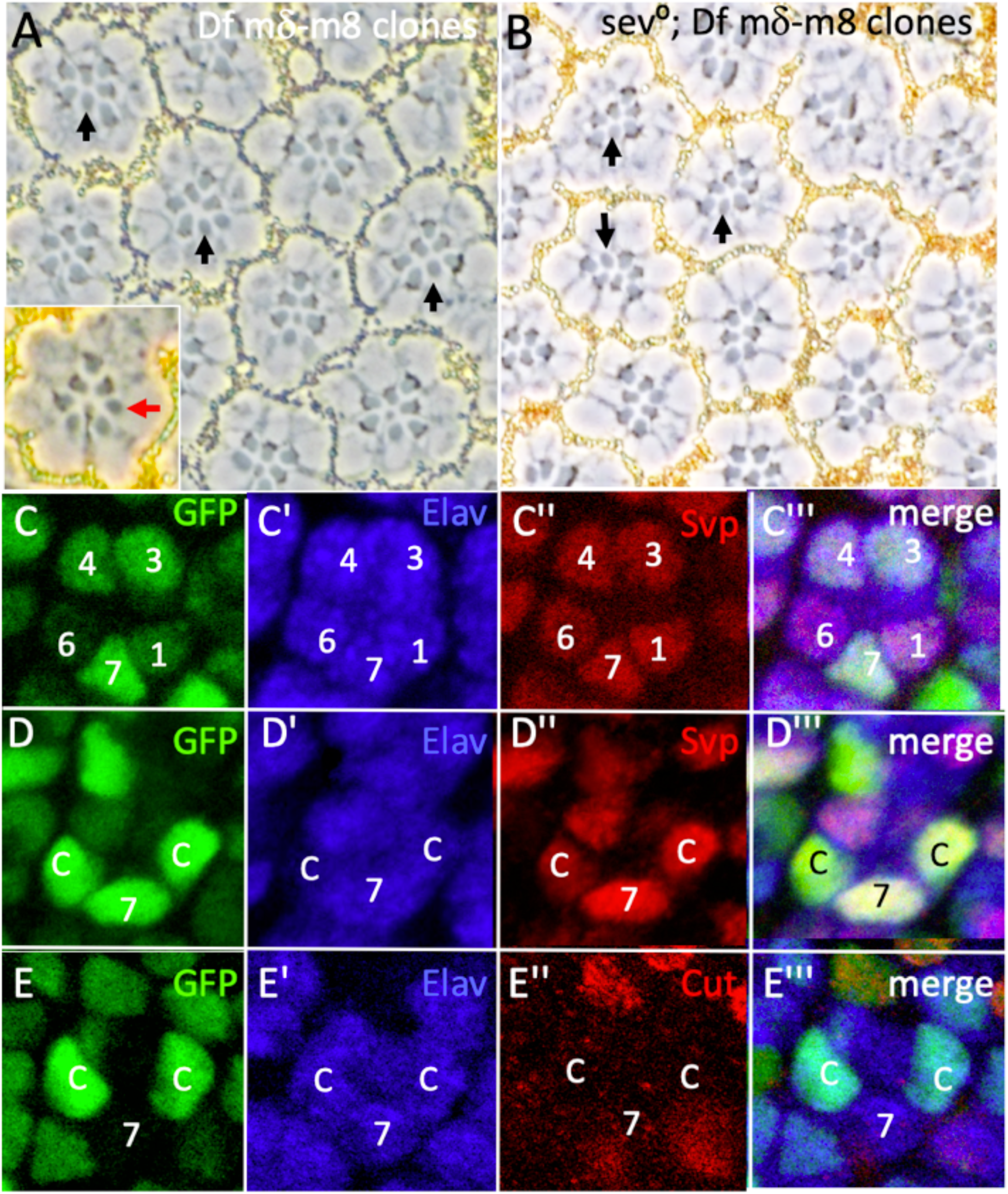
Phenotypes of *Df(3R)mδ-m8* clones. *GMR.flip* was used to generate cells homozygous for *Df(3R)mδ-m8* in R1/6/7 and cone cells precursors. (A) When cells are homozygous for the deficiency (lacking pigment) in the R7 position (arrowheads) they appear as large rhabdomere R1/6-type cells. Inset shows a mosaic ommatidium with seven large-rhabdomere cells. (B) Clones of *FRT82 Df(3R)mδ-m8* in a *sev°* background show mutant cells in the R7 position appearing as R1/6-type photoreceptors. (C-E) images from third instar discs. Mutant cells are brightly labeled in green. (C) When R7 precursors are mutant (bright green nuclei) they appear as R1/6 types as evidenced by their expression Elav (blue) and Svp (red). (D) Shows mutant cells (bright green nuclei) in R7 and two cone cells precursors. All three express Elav (blue) and Svp (red) indicating that their transformation into R1/6 cells. (E) Mutant cells (bright green) in two cone cells precursors express the photoreceptor marker Elav (blue) but not the cone cell marker Cut (red).

If the supernumerary large-rhabdomere cells resulted from cone cell precursors losing their N-induced inhibition to photoreceptor specification, we inferred that R7 precursors would similarly lose it. Since Sev is required in the R7 precursor to overcome that inhibition, then it should not need Sev when it lacks *E(spl)* gene functions. We induced *Df(3R)E(spl)mδ−m8* clones in a *sev°* background and observed that mutant cells in the R7 positions differentiated as photoreceptors (of the R1/6 class, because they also lack the R7 specifying information) (Fig.7B black arrows). Collectively, these results suggest that *E(spl)* gene functions are required for N-directed decision to become a photoreceptor rather than a cone cell, and for the choice of the R7 versus R1/6 fates.

To evaluate these suggestions, we examined the phenotypes of *E(spl)-complex* clones in the developing eye disc, again induced in R1/6/7 and cone cell precursors using *GMR.flip*. When the *Df(3R)E(spl)mδ−m8* chromosome arm is homozygous, the GFP expression from its *E(spl)mδ*.*GFP^NLS^* reporter is doubled and nuclei appear bright green compared to the weaker heterozygous expression. This provided a facile method for identifying the *E(spl)-complex* mutant cells. Whenever cells in the R7 positions were mutant (bright green), they appeared as R1/6 types – expressing Svp and Elav (Fig.7C). This comports with the adult mosaic analyses, in which cells in the R7 positions lacking the *E(spl) complex* genes appeared as R1/6 types. When cells in the cone cell positions were similarly mutant, they frequently transformed into R1/6-like cells. Fig.7D shows three mutant cells (R7 and two cone cell precursors are bright green) and each expresses Elav (blue) and Svp (red), typical of R1/6 cells. Fig.7E shows a mosaic ommatidium in which two cone cell precursors (bright green) are lacking the *E(spl)-complex* and express the photoreceptor marker Elav and not the cone cell label Cut, again highlighting the need for *E(spl)-complex* genes to prevent the specification of cone cell precursors as photoreceptors. However, not all mutant cone cells behaved in a manner that suggested a precise switches to the R1/6 fate. Rather, there was a range of phenotypes with, for example, some not displaying Cut and gaining Svp expression without the correlating expression of Elav. Such variability is evident in Fig.7D. Here the cone cell to the right displays strong Elav and Svp expression whereas the one to the left shows weaker expressions of both. This contrasts with mutant cells in the R7 positions which transform robustly into R1/6 types in the absence of the *E(spl)-complex*. Thus, we detect a difference in sensitivity between the cone cell and R7 precursors to their loss of *E(spl)* gene functions. We address this difference in the Discussion,

### The gene targets of the E(spl) transcription factors

The experiments above suggest that E(spl) transcription factors mediate two of the N actions in the R7 precursor; the choice to become a photoreceptor rather than a cone cell, and that between the R7 and R1/6 fates. In this section we investigate the target genes of E(spl) proteins in these processes.

### (i) E(spl) repression of *phyl* transcription

The choice to become a photoreceptor is mediated by the degradation or persistence of the Ttk repressor transcription factor. This in turn is controlled through regulation of *phyl* transcription, and we previously documented that N activity increased *yan* transcription^3^ and inferred that the accruing Yan protein opposed *phyl* transcription. Here, we document roles for E(spl) proteins also in repressing *phyl* transcription. We previously described the expression pattern of the *phyl.GFP^nuc^* reporter; it is expressed strongly in photoreceptor precursors but considerably less so in the presumptive cone cells^3^. When *E(spl)m8* was expressed under *sev.gal4* transcriptional control, R1/6/7 precursors frequently showed reduced expression of the *phyl.GFP^nuc^* reporter (Fig.8A; the expression levels in the R1/6/7 precursors is significantly less than their R3/4 counterparts). This reduction correlates with the transformation of the R1/6/7 precursors into cone cells as evidenced by their expression of the cone cell label Cut (red) and the absence of the photoreceptor maker Elav (blue) (Fig.8A,A”). Although there is a significant drop in *phyl.GFP^nuc^* expression in R1/6/7 precursors it does not drop to the low level of the cone cell precursors. Fig8.B shows five cone cell precursors as evidenced by their Cut expression, but the cells derived from presumptive R1/6/7s (as inferred from their positions in the cluster) show higher *phyl.GFP^nuc^* compared to the cells in the native cone cell positions. The GFP brightness in this figure has been increased to highlight the difference in expression levels between the R1/6/7 and native cone cell precursors. As result, the R1/6/7 precursors appear to show stronger staining than the that shown in Fig.8A. We also note that the Cut expression in the presumptive R1/6/7 cells is stronger than the native cone cells. These differences are revisited in the Discussion.

**Figure 8.**
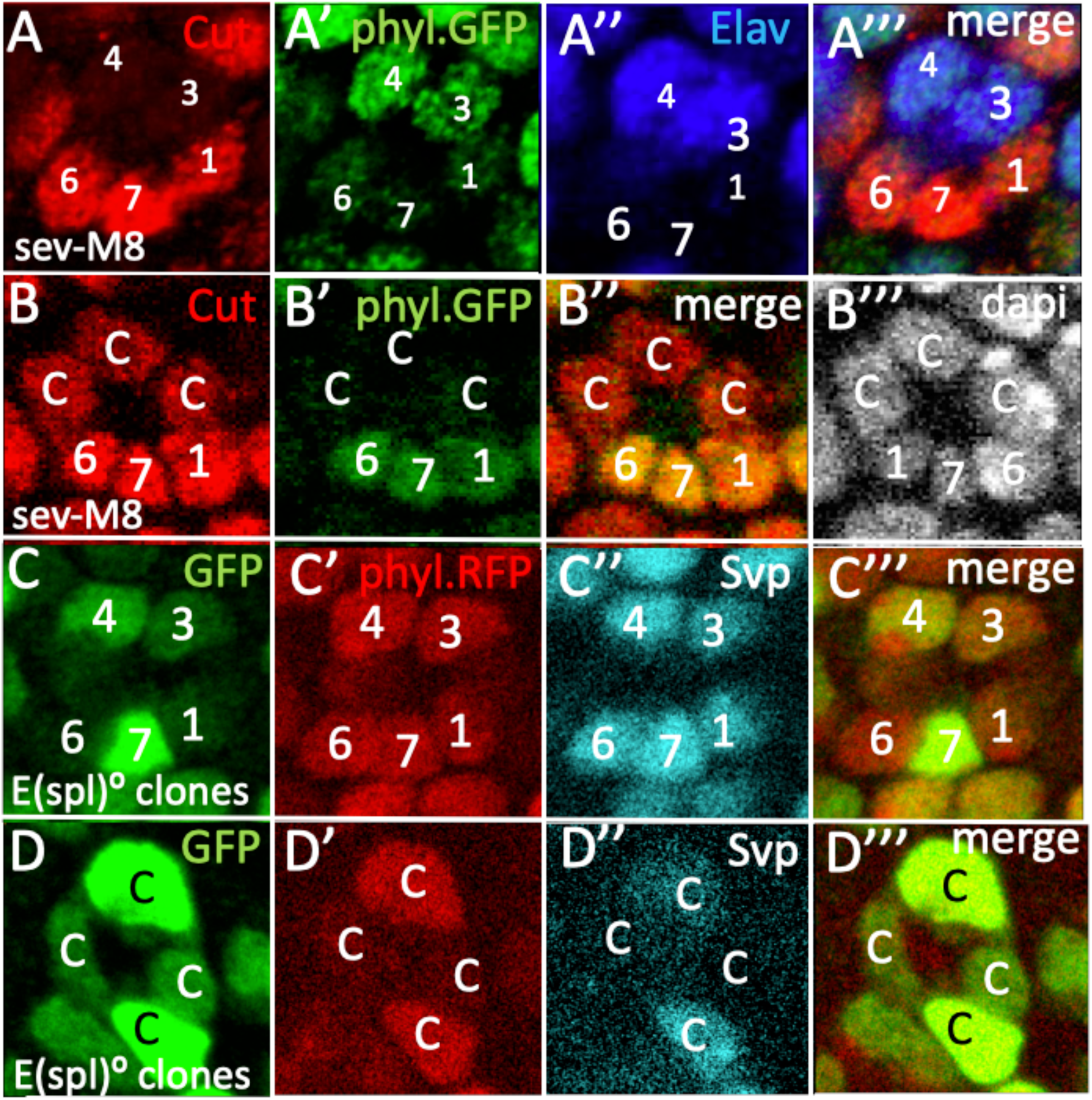
E(spl) repression of *phyl* transcription in eye discs (A,B) Effects of *UAS.E(spl)m8* on *phyl.GFP* expression. (A) When a *UAS.E(spl)m8* transgene is expressed by *sev.Gal*, R1/6/7 precursors transform into cone cells evidenced by their Cut expression and absence of Elav (A’’), and show reduced *phyl.GFP*^NLS^ expression (A’, compare expression of R1/6/7 with that of R3/4). (B) When a *UAS.E(spl)m8* transgene is expressed by *sev.Gal*, an ommatidium shows five cone cells evidenced by their expression of Cut (red). The cells in the R1/6/7 position show higher levels of *phyl.GFP*^NLS^ expression than the three native cone cells. Note, the brightness of the image in B’ has been increased to highlight the difference in expression levels between the cells in the R1/6/7 positions with native cone cells. These R1/6/7 precursors thus appear brighter than similar cells shown in A’. (C,D) Mutant clones of *Df(3R)mδ-m8* (bright green). (C) A *Df(3R)mδ-m8* mutant R7 shows normal expression of *phyl.RFP^NLS^*as it transforms into an R1/6 evidenced by its expression of Svp (silver (D) Cone cell precursors mutant for *Df(3R)mδ-m8* (bright green) show increased *phyl.RFP^NLS^*expression (red) as they transform into R1/6 cells evidenced by their Svp label (silver).

The experiments above suggested that E(spl) transcription factors can repress *phyl* transcription. If E(spl) proteins indeed normally perform this function, then the removal of the *E(spl)-complex* should lead to a de-repression *phyl* transcription in cone cell precursors. Since *Df(3R)E(spl)m8−mδ* cells are labeled by GFP we could not use *phyl.GFP^nuc^*and therefore generated a new *phyl* transcriptional reporter using *RFP^nuc^* to monitor transcription levels. This *phy.RFP^nuc^* reporter showed an identical expression pattern to *phyl.GFP^nuc^* with expression higher in the presumptive photoreceptors than the cone cell precursors (not shown). Fig. 8C shows a mosaic cluster in which R7 lacks the *E(spl)-complex* (bright green) and shows normal levels of *phy.RFP^nuc^* expression (Fig. 8C). If E(spl) proteins normally suppress *phyl* expression, then in their absence one would expect *phyl.RFP* expression levels to rise in R7 precursors. We address the reasons why this does not occur in the Discussion.

We next examined *Df(3R)E(spl)mδ−m8* clones in cone cell precursors and observed their corresponding increased expression of *phy.RFP^nuc^*; Fig.8D shows two mutant cone cell precursors (bright green) showing up-regulation of RFP^nuc^ expression when compared to their wild type counterparts. Thus, when cone cell precursors lack *E(spl)* gene functions they transform into photoreceptors and in doing so they show a correlating upregulation of *phyl* transcription. We infer therefore that genes of the *E(spl)-complex* normally function to repress *phyl* transcription.

### (ii) E(spl) repression of *svp* transcription

When R7 precursors lack *E(spl) gene* functions, they transform into R1/6 type photoreceptors defined by the expression of the Svp transcription factor. This suggests that E(spl) transcription factors suppress Svp expression, and we asked whether this is achieved through the repression of *svp* transcription. To investigate this, we examined eye discs in which *E(spl)m8* was expressed by *sev.Gal4* in the presence of a *svp.lacZ^nuc^* transcriptional reporter. Here, we observed cells in R1/6 positions showing the absence of LacZ expression, indicating that E(spl)m8 acts to suppress *svp* transcription (Fig.9A). However, the frequency of such R1/6 precursor transformations was strongly attenuated when compared to similar *E(spl)m8* expression in other genetic backgrounds, and only a few examples were observed. It is not clear why the frequency of fate transformation induced by *E(spl)m8* was reduced in the *svp.lacZ^nuc^* background.

**Figure 9.**
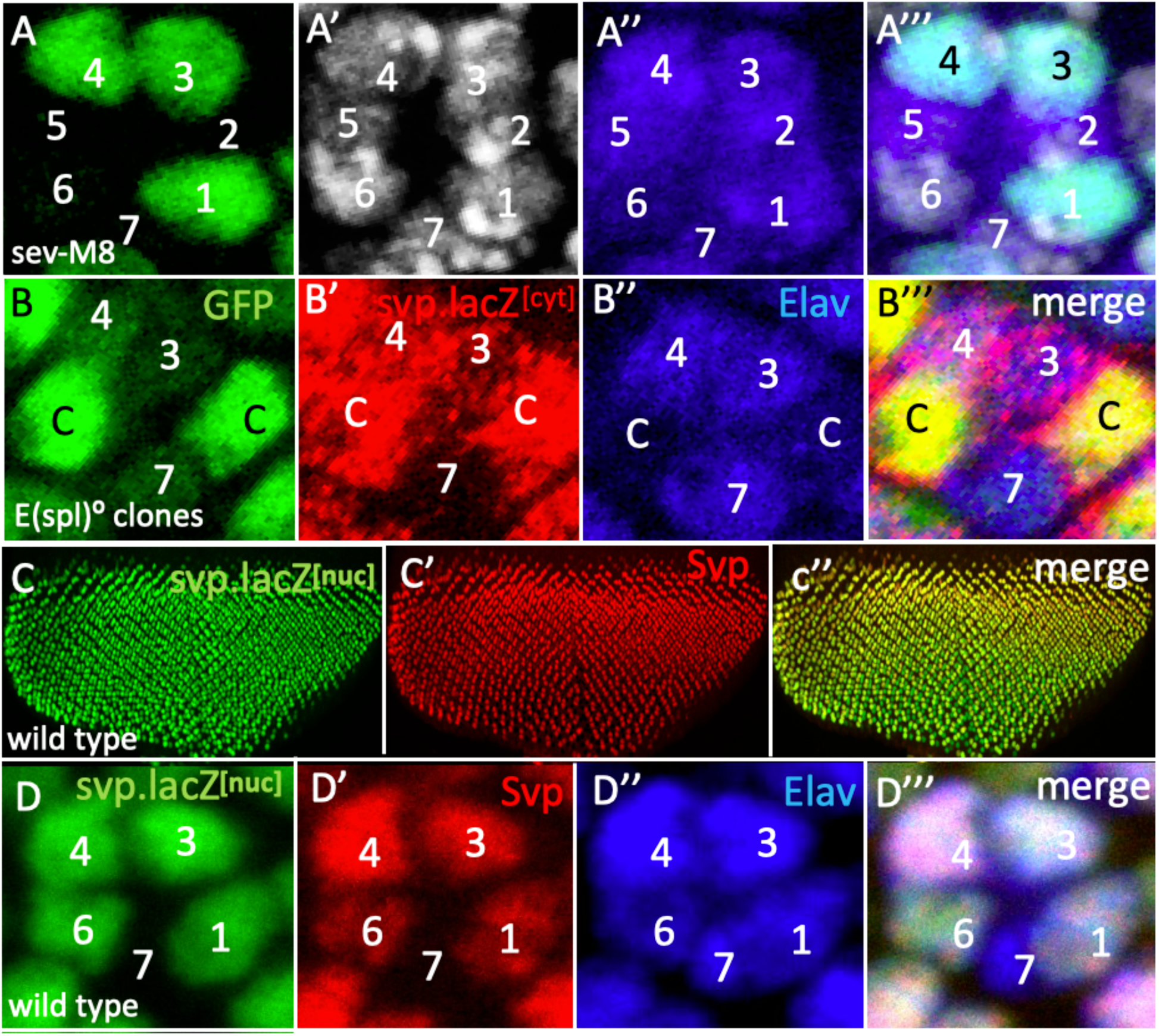
E(spl) repression of *svp* transcription in eye discs. (A) Loss of *svp.lacZ^nuc^* (green) in an R6 precursor when *UAS.E(spl)m8* is expressed by *sev.Gal4*. (B) *Df(3R)mδ-m8* mutant clones (bright green) in two presumptive cone cells show high levels of *svp.lacZ^cyt^* (red) and Elav (blue). (C) The coincident expression of svp.lacZ^cyt^ (green) and Svp protein (red) in eye discs. (D) Higher power image showing the same their coincident expression in R1/6/3/4 precursors.

If indeed E(spl) proteins repress *svp* transcription, then when the genes of the *E(spl)-complex* are removed from R7 and cone cell precursors they should display ectopic expression of a *svp* transcriptional reporter. Since the *E(spl)-complex* and *svp* lie on the right arm of the third chromosome, we used recombination to generate an *FRT82, svp.lacZ^cyt^, Df(3R)E(spl)mδ-m8* chromosome. Because we worried that the *svp.lacZ^nuc^* chromosome contained a lesion responsible for the attenuated phenotypes described above, we used a different reporter, one which displays cytoplasmic rather than nuclear LacZ expression. Clones of *FRT82, svp.lacZ^cyt^, Df(3R)E(spl)mδ-m8* were induced using *GMR.flip*, generating homozygous mutant cells labeled by their bright green expression. However, no mutant cells were detected in the R7 positions (as in the manner previously shown in Fig.7C,D) but R1/6 and cone cell precursors were liberally represented. Why no clones were detected in R7 positions is unclear and is revisited in the Discussion. However, when *E(spl) genes* were absent from cone cell precursors (bright green cells), they showed ectopic expression of the *svp.lacZ^cyt^*reporter (Fig.9B) corroborating their role in the repression of *svp* transcription. Although this cone cell result indicates the role of E(spl) genes in *svp* transcription suppression, it does not confirm that role in the R7 precursor itself. However, in eye discs, Svp protein and *svp.lacZ^nuc^*transcriptional reporter are coincidentally expressed (Fig.9A,B) with the two found in R3/4 and R1/6 precursors. This coincidence suggests that Svp protein expression is under *svp* transcription control in eye discs, and we posit that when Svp is detected ectopically in R7 precursors lacking *E(spl)* gene functions (Fig.7C,D,E), this likely results from the de-repression of *svp* transcription in those cells.

## Discussion

The R7 precursor makes two sequential fate decisions. First, it opts to become a photoreceptor rather than a cone cell, and then it decides to become an R7 rather than an R1/6 photoreceptor type. N activity plays key roles in both these decisions, including its downregulation of *phyl* transcription in opposing the specification of the photoreceptor fate, and its repression of *svp* expression to ensure that the R7 rather than R1/6 fate is specified. In this paper we find that E(spl) transcription factors mediate both these transcriptional repressions. *E(spl)* genes are classically N-responsive, and we infer that N activity in the R7 precursor achieves the repression of the *phyl* and *svp* genes by eliciting the expression of E(spl) transcription factors.

### The identity of the E(spl) transcription factors that regulate R7 specification

Our initial approach was to ectopically express E(spl) transcription factors in R1/6 precursors to identify which had the ability to re-specify these cells as R7s or cone cells. This identified three candidates, one of which one was subsequently shown not to be expressed in the R7 precursor. This highlighted the promiscuity and redundancy of the E(spl) proteins, and indicated that the ectopic expression studies were not a valid method for identifying the individual family members active in R7 specification. We therefore switched to loss of function experiments and generated a deficiency of the *E(spl)-complex*. Clones of this deficiency showed that genes of the complex are needed in R7 precursors to prevent them from being specified as R1/6 types and in presumptive cone cells to prevent them adopting a photoreceptor fate. Furthermore, we show that *E(spl)* genes are required to repress *svp* transcription in the choice to become R7, and suppress *phyl* transcription to ensure that cone cells do not transform into photoreceptors. However, there are many genes in the complex other than those that encode the E(spl) transcriptional repressors, and when we examine the phenotypes of mutant cells we cannot ascribe those phenotypes to the loss of any individual gene function or combination thereof. The ectopic studies are of value here. Since the ectopic expression of *E(spl)* genes in R1/6 precursors induced phenotypes complementary to those that the loss of the complex induced in R7 and cone cell precursors, we view this as persuasive evidence that E(spl) transcription factors indeed regulate the fate decisions made by the R7 precursor.

### The specification of the photoreceptor fate and the regulation of *phyl* transcription

The choice to become a photoreceptor versus a cone cell is regulated through the level of *phyl* transcription. If it is raised, the cell becomes a photoreceptor; if not, it becomes a cone cell. *phyl* transcription is controlled by the PntP2 and Yan ETS transcription factors. These in turn are regulated by the RTK pathway which inactivates the transcriptional repression of Yan and promotes the activation of PntP2. N activity opposes the transcription of *phyl*, and we previously documented that it induced an increase in *yan* transcription and hypothesized that the accumulating Yan proteins would titrate the ability of the RTK pathway to inactivate it. Here, we find that *phyl* transcription is also repressed by E(spl) transcription factors. We thus posit that N activity promotes the transcription of both the *yan* and *E(spl)* genes, and that the resulting proteins jointly repress expression of the *phyl* gene.

If E(spl) transcription factors indeed repress *phyl* transcription, then we would expect R7 precursors lacking the E(spl)-complex to show an increase in *phyl* expression. This we did not observe (Fig.8C). But we previously observed that *phyl* transcription did not rise in R7 precursors with abrogated N function even though that N activity represses that transcription^3^. One simple explanation is that *phyl* transcription is already maximal in the photoreceptor precursors, and removal of N activity, or the intermediaries of its functions, will not raise *phyl* expression levels further.

### The specification of the R7 fate and the repression of *svp* transcription

A key function of N activity in the R7 precursor is to prevent the expression of the *svp* gene to ensure that the cell is not specified as an R1/6 type photoreceptor. In the experiments to determine whether that N function is mediated by E(spl) transcription factors we performed both ectopic expression and loss of function analyses. Both these studies had problems. When *E(spl)m8* was expressed by *sev.Gal4* in the presence of *svp.lacZ^nuc^*, few of the phenotypes observed when *E(spl)m8* was similarly expressed in other genetic backgrounds occurred (see Fig. 8A,B). In the R1/6 precursors that were affected, there was evident reduction of expression consistent with an E(spl) role in repressing *svp* transcription (Fig.9C). We then switched to loss of function experiments and examined the effects of loss of the *E(spl)-complex* on a *svp.lacZ^cyt^* reporter. Although *E(spl)* mutant cells were frequently present in the R7 positions in experiments performed in an otherwise wild type background (Fig. 7C,D), we observed none when *svp.lacZ^cyt^* was present. Mutant cells were liberally represented in the R1/6 and cone cell precursors, and in such mutant presumptive cone cells ectopic *svp.lacZ^cyt^* expression occurred (Fig.9D), corroborating the role for *E(spl)* genes in repressing *svp* transcription. In both the ectopic expression and the loss of functional analyses, the presence of a *svp* allele (which in the case of the E(spl) clones is homozygous and genotypically mutant) appeared to influence the responses of the cells. Whether these two phenomena are related and whether they are caused by the *svp* alleles or other unrelated genetic influences is unclear. We are currently investigating these phenomena to resolve their causes. Regardless of these phenomena, there is strong evidence that E(spl) proteins indeed repress *svp* transcription and this is further bolstered by the co-expression of the Svp protein and a transcriptional reporter of its gene (Fig.9C,D). This coincidental expression suggests that Svp protein function is regulated through its gene expression, and in the many examples where we have observed ectopic Svp protein expression when E(spl) gene functions are modulated (Fig. 7C,D; Fig.8C) we infer that it results from the transcriptional activation of its gene.

### The variability of cell responses when *E(spl)* gene functions are modulated

When *E(spl)* gene functions are removed from R7 precursors they robustly transform into R1/6 type cells. But cone cell precursors showed varied responses. Sometimes they transform robustly in the manner of R7 precursors, but other times they show only weak responses (Fig.8D). We previously observed a similar phenomenon when N function was abrogated in cone cell precursors, here the cells appeared in a hybrid state expressing both Cut (the cone cell marker) and Svp (the R1/6 marker)^13^. Furthermore, when R1/6/7 precursors transform into cone cells, they show higher levels of Cut expression than the native cone cells (Fig.8B). The reasons for these variabilities are unclear, but we note that the RTK is likely more active in the photoreceptor precursors than their cone cell counterparts and this may account for differences.

### Conclusions

In the R7 precursor, N activity represses *phyl* transcription to oppose the specification of the cell as a photoreceptor. Additionally, it represses *svp* transcription to ensure the specification of the R7 rather than R1/6 type. Our work here suggests both these N functions are mediated by E(spl) transcription factors. The identities of the individual genes of the E(spl)-complex that encode these functions remain unresolved, but two such genes *E(spl)mδ* and *E(spl(m8)* are both expressed in the R7 precursor and in ectopic expression studies display the ability to perform these roles. Currently, there is no evidence that N achieves repression of the *svp* gene through anything other than E(spl) transcription factors. But N appears to engage both Yan and E(spl) transcription factors to repress expression from the *phyl* gene. Whether these two are mutually redundant or are functionally additive in their role in preventing *phyl* transcription remains unclear.

## Materials and Methods

### Generation of transcriptional reporters

For each transcriptional reporter, a homology vector was constructed containing four sequential DNA sequences. The first (5’) was genomic DNA from the gene beginning immediately 5**’** to the ATG and extending 1.5kb 5’ therefrom. The second contained the coding sequence for a fluorescent protein fused to MYC with a nuclear localization sequence followed by sv40 transcriptional termination sequences. The third contained a modified form of the *white (w)* gene coding sequence expressed under *GMR* transcriptional control (*GMR.w^+^*), flanked 3’ and 5’ either by FRTs or by LoxP sequences. Fourth was a genomic fragment beginning 3’ to the site of the most 3’ gRNA and extending 1.5kb 3’ therefrom. Three gRNAs were derived from the smallest possible region 3’ to the ATG in the first coding exon, ensuring that no non-coding sequences were deleted. gRNAs were expressed in a modified pCFD3 vector (Addgene). The homology and *gRNA* vectors were co-injected into *w^-^* embryos, and the resulting adults were crossed to *w^-^* flies. Transformants were detected by the red eyes engendered by the presence of *GMR.w^+^*. The lines were balanced, and the *GMR.w^+^*cassettes were excised using Flipase or Cre expression. Each line was checked by genomic PCR to validate its site of insertion and by complementation tests with extant mutants of the targeted gene.

### Generation of GMR.flip

The Flipase coding sequence was cloned into an attB vector immediately downstream of the GMR enhancer sequences. A flip-out cassette (>GMR.w+>) was inserted between the two, and was used as a transformation marker. The construct was transformed into attP2 at 86F after which the flip-out cassette was excised.

### Fly lines

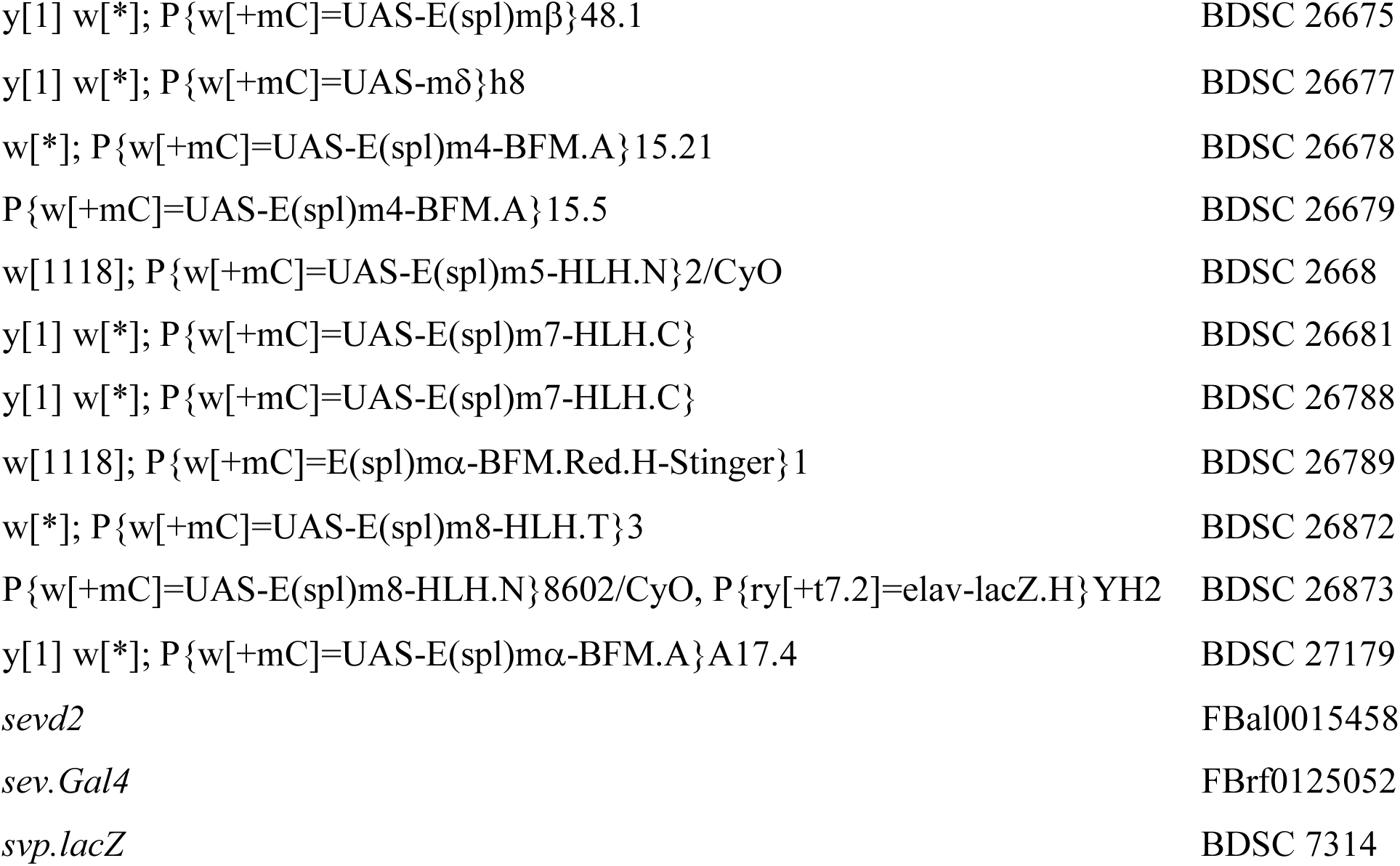

### Immunohistochemistry and histology

Protocols for adult eye sectioning^21^ and antibody staining^13^ have been described previously. All antibodies used have been previously validated, and were used in the manner described^22^

### Data validation

For adults eye experiments, the frequency of the individual phenotypes is given in the appropriate Results section. For each eye disc experiments, at least 10 developing retinas were examined, and within them, a minimum of 10 ommatidia of the appropriate stage and correct genotypic constitution were scrutinized. For example, when we examined the effects of various genetic conditions on R7 precursors, we examined at least 10 such cells to confirm the invariability of the result. But, as described in the text, the corresponding cone cell precursor phenotypes were variable and, in such situations, many more examples were scrutinized to provide an appreciation of the range of those phenotypes.

## Acknowledgments

We thank Jason Rudas for expert technical assistance with the Crispr transformations.

## Data and Resource Availability

All relevant data and resource can be found within the article.

